## Supplemental Fig. 1 for "*In vitro* one-pot construction of influenza viral genomes for virus particle synthesis based on reverse genetics system"

### Title

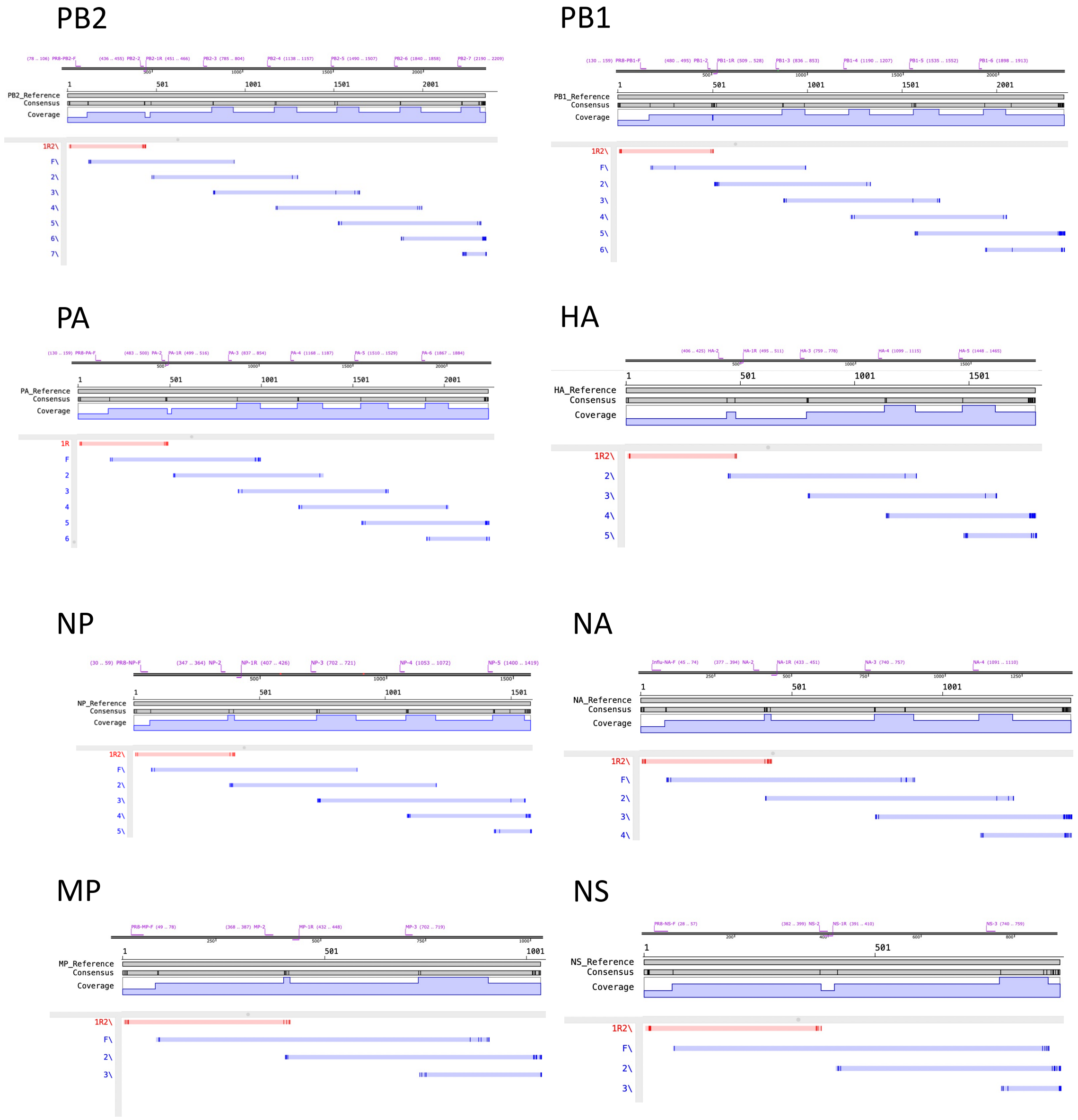


**Figure S1.** The results of Sanger sequencing of RT-PCR products from IVOC-synthesized viral genomic RNA are mapped to the reference, which is gene encoding region of the plasmids used for transfection, via GENETYX-NGS. The position of sequence primer is shown above each mapping result. The darker parts of each sequencing result indicate differences from the reference sequence. Sequencing results completely match the reference sequence except for the ends where the sequence accuracy is reduced.
